# STAT1 Promotes PD-L1 Activation and Tumor Growth in Lymphangioleiomyomatosis

**DOI:** 10.1101/2024.12.11.627871

**Authors:** Tasnim Olatoke, Erik Y. Zhang, Andrew Wagner, Quan He, Siru Li, Aristotelis Astreinidis, Francis X. McCormack, Yan Xu, Jane J. Yu

## Abstract

Lymphangioleiomyomatosis (LAM) is a cystic lung disease that primarily affects women. LAM is caused by the invasion of metastatic smooth muscle-like cells into the lung parenchyma, leading to abnormal cell proliferation, lung remodeling and progressive respiratory failure. LAM cells have TSC gene mutations, which occur sporadically or in people with Tuberous Sclerosis Complex. Although it is known that hyperactivation of the mechanistic target of rapamycin complex 1 (mTORC1) due to TSC2 gene mutations contributes to aberrant cell growth in LAM lung, tumor origin and invasive mechanism remain unclear. To determine molecular drivers responsible for aberrant LAM cell growth, we performed integrative single-cell transcriptomic analysis and predicted that STAT1 interacts with Pre-B cell leukemia transcription factor (PBX1) to regulate LAM cell survival. Here, we show activation of STAT1 and STAT3 proteins in TSC2- deficient LAM models. Fludarabine, a potent STAT1 inhibitor, induced the death of TSC2- deficient cells, increased caspase-3 cleavage, and phosphorylation of necroptosis marker RIP1. Fludarabine treatment impeded lung colonization of TSC2-deficient cells and uterine tumor progression, associated with reduced percentage of PCNA-positive cells in vivo. Interestingly, IFN-γ treatment increased STAT1 phosphorylation and PD-L1 expression, indicating that STAT1 aids TSC2-deficient tumor cells in evading immune surveillance in LAM. Our findings indicate that STAT1 signaling is critical for LAM cell survival and could be targeted to treat LAM and other mTORC1 hyperactive tumors.

## INTRODUCTION

Lymphangioleiomyomatosis (LAM) is a rare and life-threatening disease characterized by infiltration of abnormal neoplastic smooth muscle-like cells in the lung parenchyma, leading to cystic changes and progressive respiratory failure (*1, 2*). LAM demonstrates a strong gender specificity in women of reproductive age, with an estimated global prevalence of nineteen cases per million adult women (*3*). Despite knowledge that LAM is driven by mutations of tuberous sclerosis complex (TSC 1 and 2) genes in the Akt/mTORC1 pathway, mechanisms underlying LAM pathogenesis remains unclear. The current preferred treatment regimen of inhibiting mTOR is suppressive as opposed to curative, stabilizing lung function in some LAM patients but refractory in other LAM patients (*4–6*). A deeper understanding of signaling pathways dysregulated in LAM is essential to developing improved remission therapies for patients with LAM and TSC-associated disorders.

Integrative single-cell genomics analyses of LAM lung showed activation of uterine-similar HOX/PBX transcriptional network in LAM^CORE^ cells compared to other lung cells, suggesting the uterus as one of the metastatic origins of LAM cells (*7, 8*). This corroborates prior evidence of histologically similar LAM lesions in the uterus of LAM patients and female hormone responsiveness of LAM tumors (*9–11*). Although HOX genes are a family of evolutionary conserved transcription factors that are typically active during the developmental stage, reports indicate that HOX/PBX signaling is critical for female reproductive tract maintenance. Dysregulation of HOX/PBX is implicated in some female reproductive cancers (*12–14*). Prior evidence indicates nuclear colocalization of PBX1 with HOXD11, a cofactor that enhances the DNA binding specificity in LAM lung nodules and TSC2-null LAM models(*7, 8*). We recently showed that aberrant HOXD11/PBX1 activation promotes the viability and lung colonization of TSC2-null cells in our murine model (*8*). Further single-cell analysis of active transcriptional regulatory networks in the LAM lung predicted STAT1 and STAT3 as direct transcriptional targets of PBX1.

STAT proteins are critical for control of cell cycle progression, proliferation and apoptosis, and their dysregulation is associated with tumorigenesis (*15*). Overexpression of STAT1 and increased levels of phosphorylated STAT1 is associated with oncogenesis. STAT1 is overexpressed in breast (*16*) and ovarian tumors (*17*) and elevated phosphorylated STAT1 positively correlates with decreased survival rates in patients with lung cancer (*18*). Moreover, interferon-induced STAT1 phosphorylation is thought to mediate tumor immunity by increasing programmed cell death ligand 1 (PD-L1) expression in various cancers (*19–22*). LAM lung and TSC2-null immunocompetent mice have demonstrated upregulation of PD- 1/PD-L1 immune checkpoint proteins and enhanced T-cell infiltration, suggesting that LAM cells may be escaping immune detection by activating the PD -1/PD-L1 pathway (*23, 24*). We have shown that PBX1 depletion increases the expression and phosphorylation of STAT1. However, the mechanism and pathological consequence of STAT1 phosphorylation in LAM lesion growth and immune evasion remains unclear.

To examine the consequence of STAT1 activation in LAM, we analyzed integrative omics data and performed validation experiments using LAM models in vitro and in vivo to delineate mechanisms by which HOX/PBX1-induced STAT1 signaling promotes lung metastasis and tumor immunity in LAM.

## RESULTS

### 1. Integrative transcriptomics analysis of LAM lung predicts STAT1 phosphorylation in LAM Lung

Unphosphorylated STAT1 protein typically remains quiescent in the cytoplasm until they are phosphorylated and translocated to the nucleus where they activate genes involved in cell proliferation, apoptosis and survival (*25*). STAT1 phosphorylation is intricately regulated under physiologic conditions. However, aberrant STAT1 phosphorylation promotes proliferation and invasion of tumor cells (*26*). Integrative omics prediction data suggests that STAT1 gene is enriched in LAM^CORE^ cells (**Fig. 1A**). LAM lung express decreased levels of TSC2 and elevated levels of STAT1 compared to non-LAM lung **(Fig. 1B)**. To validate the activation of STAT1, we performed immunostaining of phospho-STAT1 in the LAM lung. Immunofluorescence staining showed the nuclear accumulation of phospho-STAT1 in LAM lung nodule cells as opposed to non-LAM lung cells **(Fig. 1Ci-xvi)**. Our scRNAseq analysis predicted that PBX1 and STAT1 work cooperatively to promote LAM cell survival (*8*). TSC2-null LAM patient-derived cells exhibited increased expression and phosphorylation of STAT1 and STAT3, concomitant with elevated PBX1 expression **(Fig. 1D, E)**. Conversely, treatment with HTL001, a competitive PBX1 binding antagonist (*27*), specifically decreased the protein levels of PBX1, reduced phosphorylation of STAT1, and induced phospho-RIP1, in TSC2- deficient cells, but not in TSC2-addback cells **(Fig. 1F).** These data demonstrate that PBX1- dependent activation of STAT1 is a critical to LAM cell survival.

**Figure 1:**
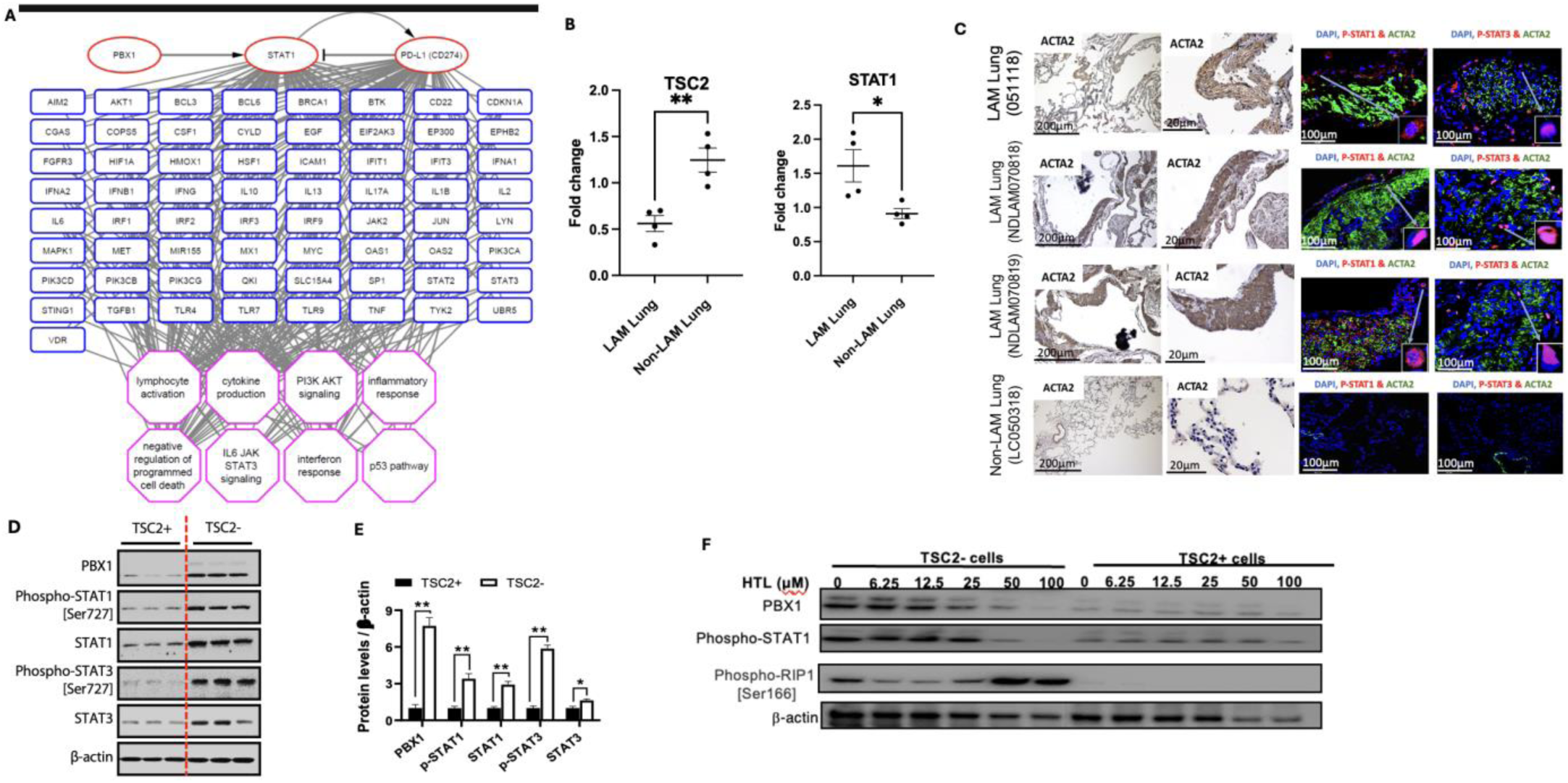
STAT1 and STAT3 are phosphorylated in LAM Lung. (**A**) Transcript levels of STAT1 and TSC2 in LAM Lung and Non-LAM lung using reverse transcription quantitative polymerase chain reaction (RT-qPCR). (**B**) LAM lung tissues and a non-LAM lung tissue were stained for ACTA2 to show nodular lesions. Immunofluorescence staining shows nuclear localization of phospho-STAT1 and phospho-STAT3 in LAM lung nodules. **(C)** Immunoblotting was done to assess protein levels of PBX1, STAT1, STAT3, phospho-STAT1 and phospho-STAT3 in TSC2- and TSC2+ LAM patient-derived cells. (**D**) Densitometry analysis of immunoblots (**E**) TSC2- and TSC2+ cells treated with 0-100uM HTL001 (a competitive PBX1 binding antagonist). Immunoblot analysis of PBX1, phospho-STAT1 and phospho-RIP1 levels; β- actin as a loading control.

### 2. TSC2 negatively regulates the activation of STAT1 and STAT3

We have identified and validated STAT1 and STAT3 activation in LAM Lung nodule cells. To define the molecular mechanisms responsible for STAT1 and STAT3 activation, we assessed their expression pattern in other TSC2-null models. Phospho-STAT1 was prominent in the nucleus of TSC2-null 621-101 cells, but not in TSC2-addback 621-103 cells **(Fig. 2A)**. Similarly, phospho-STAT3 was more abundant in the nucleus of TSC2-null cells relative to TSC2-addback cells **(Fig. 2B)**. Next, we examined LAM uterine tissues and identified desmoplastic uterine lesions, consistent with malignant neoplasms (*28, 29*). LAM uterine cells exhibited prominent nuclear accumulation of phospho-STAT1 and phospho-STAT3 compared to non-LAM lesions **(Fig. 2C)**. Moreover, we found nuclear accumulation of phospho-STAT1 and phospho-STAT3 in myometrial tumor cells from Tsc2KO compared to wildtype mice **(Fig. 2D)**. We also found positive immunoreactivity of phospho-STAT1 in lung metastatic cells from xenograft tumors of Tsc2-null ELT3 cells, but not in Tsc2-addback ELT3-T3 cells **(Fig. 2E)**. Together, our data implies that TSC2 negatively regulates STAT1 and STAT3 activation.

**Figure 2:**
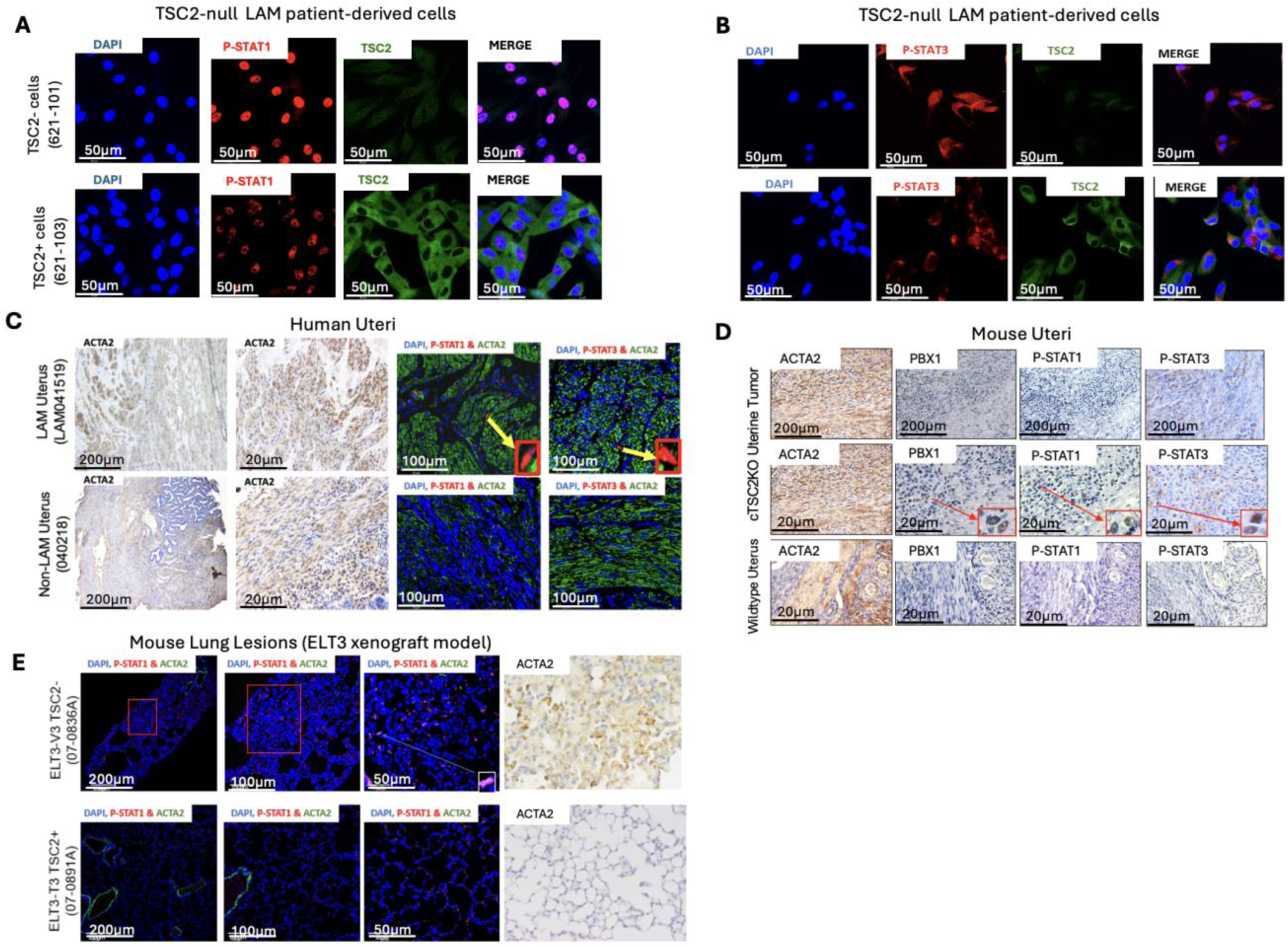
TSC2 negatively regulates the activation of STAT1 and STAT3. (**A**) Immunofluorescence staining of TSC2 and phospho-STAT1 in TSC2- (621–101) and TSC2+ (621–103) cells. Confocal staining shows nuclear staining of phospho-STAT1 in TSC2-null cells. (**B**) Immunofluorescence staining of TSC2 and phospho-STAT3 in (621–101) and TSC2+ (621–103) cells. (**C**) Formalin-fixed paraffin embedded (FFPE) uterine tissues from a LAM patient and a non-LAM patient collected during hysterectomy. LAM nodular lesions indicated by smooth muscle actin (ACTA2). Immunofluorescence staining of phospho-STAT1 and phospho-STAT3. (D) FFPE uterine tissues from uterine-specific TSC2 knockout mice with enlarged uteri and from wildtype mice. Representative images of immunohistochemical staining of phospho-STAT1 and phospho--STAT3. (**E**) Tsc2-null ELT3-V3 or TSC2-addback ELT3-T3 cells (2 × 10^6^) were subcutaneously injected into both flanks of 8-week-old female CB17/C.B-Igh-1^b^/IcrTac-Prkdc^scid^ (SCID) mice, resulting into micro metastasis of TSC2-null lesions. Representative images of immunofluorescence staining of phospho-STAT1 in xenograft tumors.

### 3. STAT1 suppression by Fludarabine triggers cell death in TSC2-null cells

Our previous network analysis of LAM multiome predicted that PBX1 and STAT1 share target genes associated with “negative regulation of cell death” in LAM^CORE^ cells (*8*), suggesting that aberrant STAT1 activation induces the survival of LAM cells. TSC2-null cells have constitutive activation of mTORC1, which is suppressed by Rapamycin. However, Rapamycin treatment did not affect the phosphorylation of STAT1 in TSC2-null (621–101) cells compared to TSC2-addback (621–103) cells **(Fig. 3A)**, suggesting an alternative pro-survival pathway for mTOR hyperactive cells. Combination therapy has been demonstrated to increase curative efficacy of tumor cells compared to monotherapy (*30*). To determine the potential efficacy of combined suppression of STAT1 and mTORC1 on cell survival in vitro, we treated LAM patient-derived cells with Fludarabine singly or in combination with Rapamycin. Combinatorial treatment of Fludarabine and 20 nM Rapamycin decreased the viability of TSC2-null 621-101 cells (GI_50_=18.3 μM) (blue circle), compared to TSC2-expressing 621-103 cells (GI_50_=117 μM) (red square) within 72 hours **(Fig. 3B)**. Interestingly, the concentration of Fludarabine (GI_50_= 31.1 μM) (black circle) needed to reduce the viability of cells by 50% was almost double the concentration of Fludarabine with Rapamycin (GI_50_=18.3 μM) (blue circle) to exert the same effect in 621-101 cells **(Fig. 3B)**. Moreover, Fludarabine treatment for 48 h increased the levels of cleaved-caspase 3, phospho-RIP1, and PD-L1 in TSC2-null cells **(Fig. 3C)**. Flow cytometry assay showed that Fludarabine treatment significantly increased annexin V-positive and PI-positive TSC2-null cells, thereby reducing viable TSC2-null cells relative to TSC2-addback cells **(Fig. 3D-F)**. Collectively, our result suggests that STAT1 promotes the survival of TSC2-null cells in vitro.

**Figure 3:**
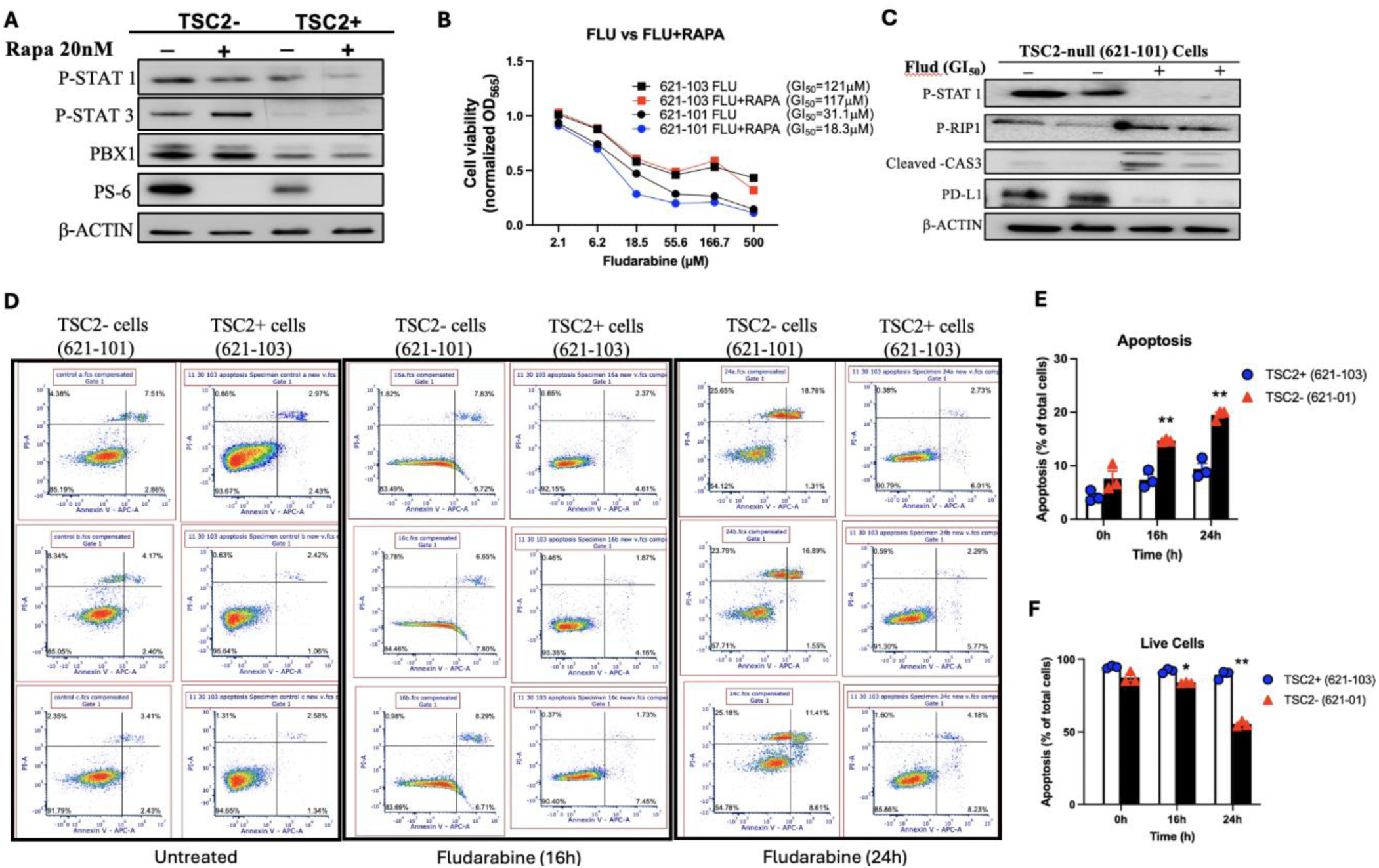
Fludarabine induces death of TSC2-null LAM-derived cells in vitro. (**A**) 621-101 cells were treated with 20nM Rapamycin. Immunoblotting analysis of PBX1, phospho-STAT1 (Ser727), phospho-STAT3 (Ser727), and phosphor-S6; β-actin as a loading control. (**B**) Viability of 621-101 and 621-103 cells was assessed 72 hours after treatment with Fludarabine alone and Fludarabine with 20nM Rapamycin. Black square, 621-103 Fludarabine (GI_50_=121 µM); Red square, 621-103 Fludarabine and 20nM Rapamycin (GI_50_=117 µM); Black circle, 621-101 Fludarabine (GI_50_=31 µM); Blue circle= 621-101 Fludarabine and 20nM Rapamycin (GI_50_=18.3 µM). GI50 was calculated using CompuSyn. (**C**) Immunoblot analysis of phospho-STAT1 and cell death markers phospho-RIP1 and Cleaved-caspase 3 in Fludarabine-treated 621-101 cells. β- actin as a loading control. (**D**) 621-101 (TSC2-) and 621-103 (TSC2+) cells were treated with xx Fludarabine for 16 and 24 h, harvested, then stained with Annexin V: FITC Apoptosis Detection Kit. Cell death was analyzed by flow cytometry (n = 3). (**E**) The percentage of apoptotic (annexin V+PI positive) and (F) viable (annexin V-PI negative) cells in total cell number was determined.

### 4. Suppression of STAT1 by Fludarabine attenuates Tsc2-null cell lung colonization and myometrial tumor growth

To examine the preclinical efficacy of blocking STAT1 activation in vivo, we first established lung colonization of Tsc2-null cells (*11*). Fludarabine treatment for 72 h did not affect lung colonization of Tsc2-null ERL4 cells in vivo (*8*). In the present study, Fludarabine potently reduced lung lesion formation by 90% twelve weeks post-cell injection (**Fig. 4A-B**). Next, we treated Tsc2KO mice with Fludarabine for six weeks. Fludarabine treatment lowered body weights and uterine weights, relative to untreated Tsc2KO and wildtype control mice **(Fig 4C-4E).** As expected, Tsc2KO mice have no TSC2 protein expression and increased phosphorylation of S6 protein. Importantly, Fludarabine treatment reduced expression and phosphorylation of STAT1 compared to untreated Tsc2KO mice, indicative of the efficiency of the treatment **(Fig 4F)**. This was associated with reduced immunoreactivity of PCNA, a tumor proliferation marker in Fludarabine treated mice relative to untreated mice **(Fig 4G, H)**. Overall, our data demonstrates that STAT1 activation promotes TSC2-null cell survival in vivo.

**Figure 4:**
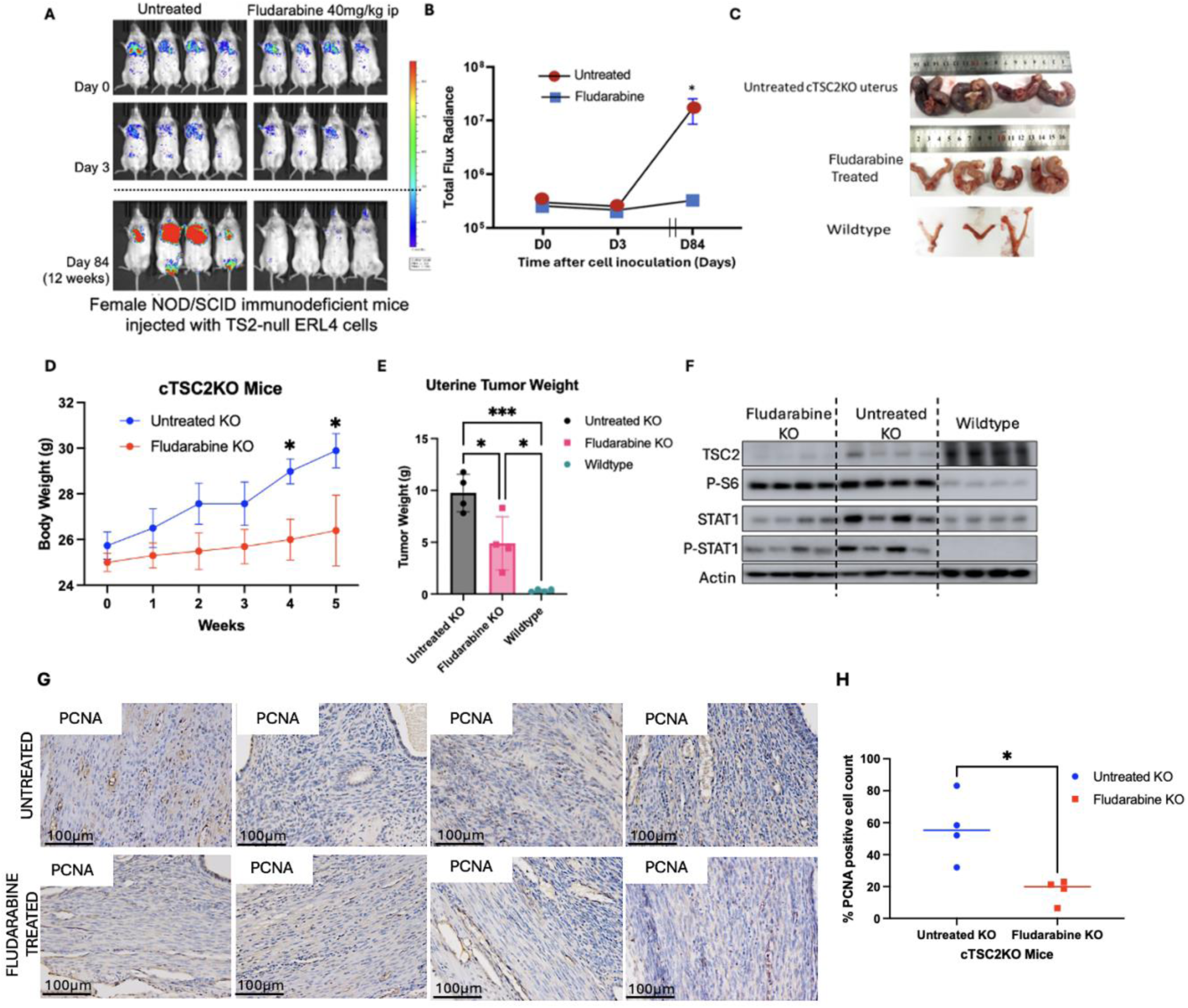
Fludarabine attenuates lung colonization and uterine tumor growth in TSC2- deficient models. (**A**) Female NSG mice were pretreated with Fludarabine (40 mg/kg ip) and intravenously inoculated with 2 × 10^5^ ERL4 cells. Mice were post-treated with Fludarabine for 72h. Bioluminescence imaging was performed for 12 weeks. (**B**) Bioluminescence signals from luciferase expressing ERL4 cells were quantified. Total photon flux per second present in the chest regions were quantified (n=4/group). (**C**) Mice with uterine-specific TSC2 deletion (cTSC2KO) were treated with Fludarabine for 6 weeks (n = 4/group). (**D**) Body weights were recorded weekly Uterine tumor weights were measured at treatment endpoint (**F**) Immunoblot of TSC2, phospho-S6, STAT1, and phospho-STAT1 in uteri from wildtype, Tsc2KO, and Fludarabine-treated mice (n=4/group). β-Actin as a loading control. (**G**) Uterine sections from Fludarabine-treated or untreated Tsc2KO were stained with PCNA. Scale bars are 100 μm. (**H**) Percentage of cells with nuclear immunoreactivity for PCNA was estimated *P < 0.05, **P < 0.01.

### 5. IFN- γ activates Phospho-STAT1/IRF1/PD-L1 pathway in TSC2-null cells

Since PD-L1 was enriched as a STAT1 target gene and interferon response **(Fig 1A)**, we sought to determine the relevance of PD-L1 in LAM pathogenesis. We stained LAM lung tissues and observed strong immunoreactivity of PD-L1 in LAM lung nodule cells **(Fig. 5A)**. PD-L1 accumulation was also abundant in myometrial cells from Tsc2KO mice relative to control mice **(Fig 5B)**. IFN-γ-mediated STAT1 activation promotes immune evasion by stimulating PD-L1 expression in various tumors (*19–22*). Upon stimulation with IFN-γ, STAT1 is activated via phosphorylation, which induces IRF1 expression. IRF1 binds to PD-L1 promoter region to induce PD-L1 expression and membrane translocation (*31*). To determine whether STAT1 phosphorylation mimics this effect in TSC2-null cells, we treated 621-101 cells with IFN-γ for 48 h and observed prominent cytoplasmic localization of PD-L1 in response to IFN-γ **(Fig. 5C)**. IFN-γ potently increased the protein levels of STAT1, IRF1, PD-L1, and phosphorylation of STAT1 in TSC2-null cells **(Fig. 5D-E)**. IFN-γ promotes the migration of TSC2-null cells in a scratch wound-healing assay **(Fig. 5F)**. Our data reveals that IFN-γ is critical in promoting tumor growth in TSC2-null cells in vitro.

**Figure 5:**
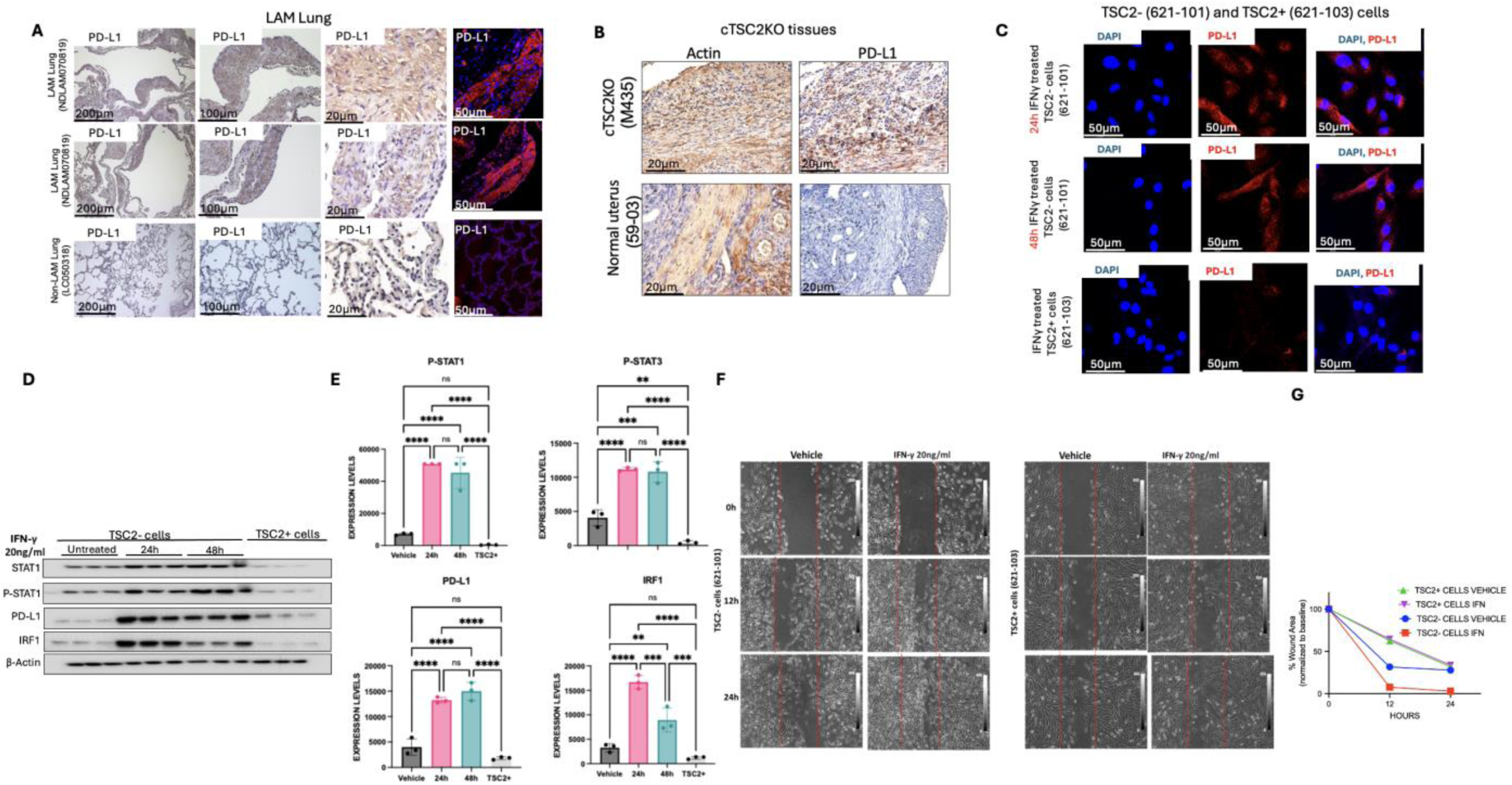
IFN- γ activates PSTAT1/IRF1/PD-L1 pathway in TSC2-null cells. (**A**) Immunohistochemical staining of ACTA2 and PD-L1 in LAM lungs. Representative images of 2 cases are shown. Scale bars, 200 μm (a, e, i), 100 μm (b, f, j), 20 μm (c, g, k) and 50 μm (d, h, l). (**B**) Immunohistochemical staining of ACTA2 and PD-L1 in uterine tissues from Tsc2KO and wildtype mice. Scale bars are 20 μm. (**C**) TSC2-null and TSC2-addback LAM patient-derived cells were treated with IFN-γ (20ng/ml) for 48 h. Immunofluorescence staining of PD-L1. Scale bars are 50 μm. **(D**) TSC2-null LAM patient-derived cells were treated with 20 ng/ml IFN-γ for 24 h and 48 h. Immunoblot analysis of STAT1, phospho-STAT1, PD-L1, and IRF1 (n=3). β-actin as a loading control. **(E)** Densitometry of STAT1, PSTAT1, PDL1 and IRF1 blots (n=3). *P < 0.05, **P < 0.01. (**F**) Scratch Wound-Healing assay showing migration of TSC2-null and TSC2- addback LAM patient derived cells with 16h and 24h IFN-γ.

## DISCUSSION

The biggest challenge to finding curative therapy for LAM tumor cells is its elusive primary origin and its undetermined invasive mechanisms. We have previously shown compelling evidence that the uterus is likely one of the major origins of LAM cells (*7*). LAM lung and uterine cells have comparable biallelic TSC2 mutations, morphological features, and transcriptional profiles, supporting this claim (*2*). Our analysis of the LAM lung multimode data predicted an increased expression and activation of uterine-similar HOX/PBX transcriptional signaling pathway(*8*). HOXD11 and PBX1 are in the top 1% of active transcription networks in LAM^CORE^ cells. We validated HOXD11 and PBX1 activation in LAM lungs, LAM uteri and mouse models of LAM. While targeting HOXD11 and PBX1 activation in LAM has been promising, our transcription factor study of PBX1 indicates a strong affinity for STAT1 and STAT3. Collectively, this study provides evidence of the crucial role of STAT1 and STAT3 in promoting LAM cell survival.

We observed that STAT1 and STAT3 are phosphorylated in TSC2-deficient cells, with prominent nuclear accumulation in LAM lung nodules, LAM uterus, LAM-patient derived TSC2 cells and TSC2-deficient lung lesion and uterine mouse models. Although all TSC2- deficient cells show mTORC1 hyperactivity, the inhibition of mTORC1 using rapamycin did not affect activation of STAT1 and STAT3. Fludarabine is a purine analogue that reduces mRNA and protein levels of STAT1 (*32*) (*33*) (*34*), inducing apoptosis in TSC2-deficient cells. We reasoned that combinatorial inhibition of mTORC1 and STAT1 will show greater therapeutic efficacy than STAT1 or mTORC1 monotherapy. Our findings demonstrate that the dual treatment of Fludarabine and Rapamycin induced more cell death than Fludarabine alone, suggesting a potential synergism between STAT1 and mTORC1 in LAM cells.

Our lung colonization mouse model mimics the metastatic mechanism of TSC2-null in humans. We previously reported that short-term treatment of Fludarabine does not affect TSC2- lung colonization in mice (*8*). However, our long-term assessment shows a 90% decline in lung lesion burden, indicating prolonged benefits of Fludarabine. The strongest evidence for the origin of LAM cells is the uterus. Fludarabine effectively shrunk uterine tumor size in mice by reducing the rate of tumor proliferation. Other studies showed that purine nucleotides are critical for cell proliferation (*35*) (*36*). Mizoribine, an IMPDH inhibitor, induces the death of TSC2-null cells in vitro and in vivo (*37*). Future studies will warrant the specificity and on-target effect of STAT1 inhibition in LAM models.

The regulation of anti-tumor immune responses through cancer immunotherapy is one of the most innovative approaches to treating tumors. This groundbreaking approach eliminates the concerns for detrimental side effects of standard treatment strategies such as radiation and chemotherapy by leveraging the immune system to eliminate tumor cells (*38*). Tumor cells evade immune detection and elimination via several pathways, including the upregulation of immune checkpoint molecules such as programmed death ligand-1 (PD-L1). Activated T cells expressing programed death-1 (PD-1) bind to a ligand produced by tumor cells (PD-L1), resulting in T cell suppression and unchecked tumor growth. Cancer immunotherapy targets PD-1/PD-L1 interaction to restore T cell function on tumor elimination. In the present study, we show upregulation of PD-L1 in LAM lung nodule cells and TSC2-null lesion cells, consistent with previous reports (*23, 24*). We also show that Fludarabine treatment suppresses the protein levels of STAT1, concomitant with reduced protein levels of PD-L1, indicating that STAT1 regulates PD-L1 expression in TSC2-deficient tumor cells. Moreover, STAT1- mediated PD-L1 expression is under the influence of IFN-γ in the present study. Our findings show that IFN-γ promotes the proliferation and migration of TSC2-deficient cells in vitro, suggesting that STAT1 contributes to tumor immunity of LAM cells via IFN-γ regulated STAT1/PD-L1 expression. This is consistent with prior findings that STAT1 activation increased expression of PD-L1 in ovarian, melanoma, breast cancer, cervical and oral squamous cancer cell lines (*19–22*).

Our current study provides evidence for the crucial role of STAT1 activation in LAM progression. STAT1 promotes tumor proliferation by enhancing TSC2-null tumor growth and immune evasion via PD-L1. We provide a proof of concept for the potential therapeutic benefit of targeting STAT1 and/or PD-L1 in LAM and other mTORC1 hyperactive tumor disorders.

## MATERIALS AND METHODS

### Cell culture and reagents

Eker rat uterine leiomyoma-derived cells (ELT3) cells (*39*) were generously donated by Dr. C. Walker, Institute of Biosciences and Technology Texas A & M University, Houston, TX. LAM patient-associated angiomyolipoma-derived (621-101 and 621-103) and luciferase-expressing ELT3 cells were generously donated by Dr. E.P Henske, Brigham and Women’s Hospital-Harvard Medical School, Boston, MA (*11, 40*). Cells were grown in Dulbecco’s modified Eagle’s medium (DMEM) supplemented with 10% fetal bovine serum (FBS), and 1% penicillin-streptomycin-amphotericin B (PSA) at 37°C in a humidified chamber with 5% CO2. Cells were subjected to Mycoplasma testing using MycoAlert Mycoplasma Detection Kit (Lonza, #LT07) prior to experiments. Fludarabine (0-500 µM; #118218 Selleck Chemicals LLC, TX, USA), Rapamycin (20nM; Enzo #BML-A275-0025) and HTL001 (0 to 100 μM; Bio-Synthesis Inc., Lewisville, TX, USA) were used as indicated.

### Immunofluorescence staining and Confocal Microscopy

Paraffin-embedded tissue sections were dewaxed, rehydrated, and unmasked by heat-induced antigen-retrieval. Cells were cultured in a 6-well glass bottom plates and fixed with 4%PFA and permeabilized with 0.2% Triton X-100. Blocking was done in 10% goat serum (in PBST), followed by incubation with primary antibody (1:100 in 10% goat serum) at 4° C overnight, and incubation with secondary antibody (1:1,000, anti-Rabbit #A11011 and anti-mouse #A11001 Invitrogen) at room temperature. Counterstaining was done using 4’,6-diamidino-2-phenylindole (Invitrogen, #D1306). All Images were captured using a Leica Stellaris 8 Confocal Microscope and LAS X software (Leica Microsystems, Wetzlar, Hesse, Germany).

### Immunoblotting

Cells were lysed in T-PER or M-PER buffer (Thermo Fisher Scientific) containing protease and phosphatase inhibitors. Equal amounts of protein were separated using 4–20% Mini-PROTEAN TGX Precast gels (Bio-Rad) and transferred using a wet electrophoretic transfer unit (Bio-Rad) onto PVDF membranes (Millipore Sigma). Membranes were blocked in either 5% BSA or 5% non-fat dry milk and incubated with primary antibodies against Phospho-STAT1 (ser727) (#9177), STAT1 (#14994), Phospho-STAT3 (ser727) (#9145), STAT3 (#30835), PBX1 (#4342), cleaved caspase-3 (#9661), Phospho-S6 (ser235/236) (#4858), Phospho-RIP1 (ser166) (#44590), IRF1 (#8478), TSC2 (#4308) and β-actin (#3700) from Cell Signaling Technology, and PD-L1 (#GTX104763) from GeneTex. Blots were incubated in HRP-linked secondary antibodies and developed using SuperSignal West Pico PLUS Chemiluminescent Substrate (#PI34580 Thermo Fisher Scientific).

### Immunohistochemistry

Deparaffinized sections were incubated in target primary antibodies, followed by secondary antibody incubation using Mouse/Rabbit Specific HRP/DAB IHC Detection Kit (Abcam, #ab64264). Chromogenic detection of horseradish peroxidase (HRP) activity was done using DAB substrate (ThermoFisher Scientific #34002) and counterstained with hematoxylin solution (Fisher Scientific #220-100). Images were captured using phase-contrast microscopy. PCNA expression was detected immunohistochemically (CST #13110). The PCNA labeling index was assessed by counting PCNA positive stained nuclei using FIJI software.

### Cell Viability Assay

Cells were seeded at a density of 2×10^3^/well in 96-well plates and incubated for 24 h prior to treatment with inhibitors (Fludarabine 0-500 µM; Rapamycin 20nM) or vehicle for 72 h. Cell viability was determined using MTT assay. 25 µL of MTT solution was added to each well, followed by 4 h incubation of plates at 37°C. The resulting formazan crystals were solubilized using 0.04N HCl in isopropanol, followed by 1.5 h incubation at 37°C. Cell viability was measured by assessing absorbance at 560nm with a reference of 650nm. Growth inhibitory power was analyzed using CompuSyn Software.

### Flow Cytometry

Cells were treated with Fludarabine for 0, 16 h and 24h. Apoptosis was determined by staining cells with Annexin V-FITC/PI Apoptosis Detection Kit (Biolegend, #640914) according to manufacturer’s protocol. Cells were analyzed on a Attune Flow Cytometer using FCS Express Software.

### Animal studies

For lung colonization studies, luciferase expressing ELT3 cells (2×10^5^ cells) were intravenously injected into female SCID mice (Taconic Biosciences, Germantown, NY) as previously described (*11*). Animal health was monitored daily. Mice were treated with Fludarabine (40 mg kg/day, ip), or vehicle for 48h before cell inoculation and 24h after cell injection. Lung lesion development was tracked for 12 weeks.

For genetic Tsc2 deletion studies, we generated mice with uterine-specific Tsc2 deletion as previously described (*41*). At 18 weeks old, mice were randomly assigned to two groups. Fludarabine (40mg/kg) and Vehicle control group (n=4). Treatment was delivered intraperitoneally to mice five times a week for 6 weeks. Animal health was monitored daily. Animals were euthanized at study endpoint and uteri and lungs tissues were removed.

### Bioluminescent reporter imaging

Lung colonization was tracked in mice injected with luciferase expressing ELT3 cells. D-luciferin was administered to mice 10 mins before imaging and bioluminescent signals from chest regions were obtained using the Xenogen IVIS Spectrum System. Total photon flux was determined as previously described (*11*).

### Human subjects and tissue collection

Lung and uterine tissues were obtained from the National Disease Research Interchange (NDRI).

### Quantitative RT-PCR

RNA was isolated using GeneJET RNA Purification Kit (Thermo Scientific) and reverse transcribed to cDNA using High-Capacity cDNA Reverse Transcription Kit (Invitrogen). Gene expression was quantified using the Quant Studio Real-Time PCR systems (Applied Biosystems). RNA was normalized to standard control. Primers used are listed in supplementary table.

### CRISPR Knockout

CRISPR/Cas9 was employed to knockout STAT1 in TSC2-null ERL4 cells. Three sgRNA lentiviral targets (81 GGTACTGTCTGATTTCCATG, 164 GAGCAGGTCATGGAAGCGGA and 309 GATCATC CAC AACTGTCTGA) were used to guide Cas9 to cause frameshift mutations and STAT1 gene knockout. A Cas9-absent sgRNA was used as a control. ERL4 cells were transduced with CRISPR lentiviruses overnight at 37°C with 5% CO2 incubator and selected with puromycin. The first construct was the most efficient and used for animal studies. pLenti-U6-sgRNA-SFFV-Cas9-2A-Puro STAT1 construct, and control construct were synthesized by Applied Biological Materials.

### Transcriptional regulatory networks analysis of Multiome using paired gene expression and chromatin accessibility data

TRN analysis was conducted using integrative RNA-seq (n = 4) and ATAC-seq (n = 3) of the LAM^CORE^ cells. The PECA 2.0 statistical model (*42*) was used to determine gene regulatory networks. Networks were visualized using the Cytoscape software (*43*). Network display edge regulation score cutoff was in the 99th percentile, while the functional enrichment analysis was in the 90th percentile.

### Scratch Wound-Healing Assay

TSC2-null (621–101) and TSC2-addback (621–103) cells were seeded in 6-well plates (1.0 × 10^5^ cells, n=3/well) for 24h. A wound was created using a sterile pipette tip. The detached cells were washed using PBS and cells were stimulated by IFN-γ. Wound healing images were captured after 12 h and 24 h post IFN-γ stimulation using an inverted microscope. The area of wound was estimated using Image J software.

### Statistical analysis

All data in graphs are shown as mean ± SEM. Analysis and graphing were performed using Prism 10 (GraphPad software, LLC). The two-tailed Student’s *t* test was used to compare two groups and analysis of variance (ANOVA) test (Dunnett’s multiple comparisons test when comparing multiple groups with control group and Tukey’s multiple comparisons test when making multiple pairwise comparisons between different groups) for multiple group comparisons. Statistical significance was defined as *P* < 0.05.

### Study approval

The University of Cincinnati Institutional Animal Care and Use Committee (IACUC) approved all procedures described according to standards set forth in the Guide for the Care and Use of Laboratory Animals (IACUC, no. 24-06-10-01). The Institutional Review Board (IRB) of the University of Cincinnati approved all studies using de-identified clinical specimens (IRB, no. 2016-7095).

**Table 1.**
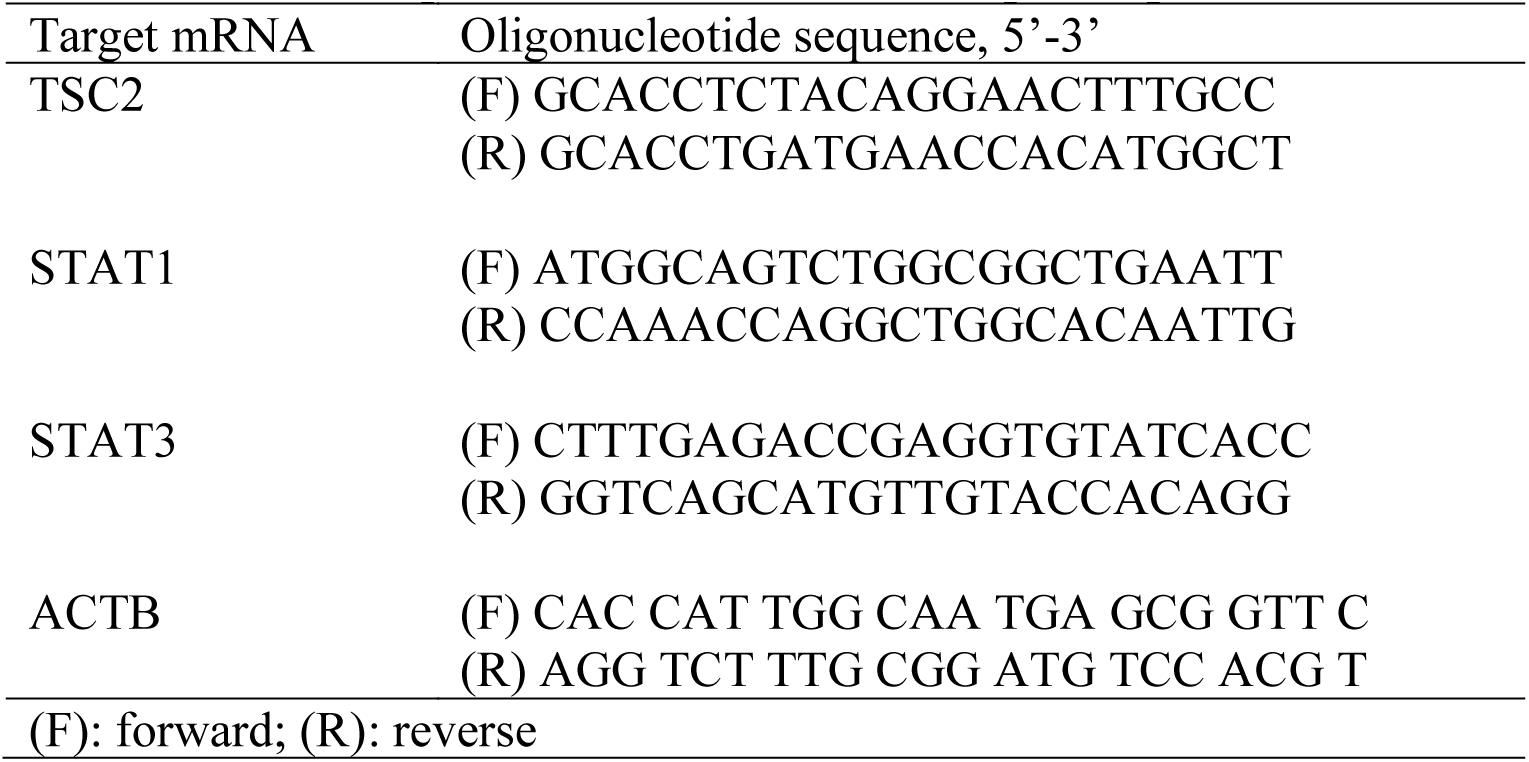
Primer sequences for reverse transcription-quantitative PCR.

## Acknowledgments

We thank The LAM Foundation and the National Diseases Research Interchange for assistance with tissue collection.

