## Supplemental Table 1 for "STAT1 Promotes PD-L1 Activation and Tumor Growth in Lymphangioleiomyomatosis"

**Table 1. Primer sequences for reverse transcription-quantitative PCR.**

| Target mRNA | Oligonucleotide sequence, 5’-3’ |
| --- | --- |
| TSC2 | (F) GCACCTCTACAGGAACTTTGCC  (R) GCACCTGATGAACCACATGGCT |
| STAT1 | (F) ATGGCAGTCTGGCGGCTGAATT  (R) CCAAACCAGGCTGGCACAATTG |
| STAT3 | (F) CTTTGAGACCGAGGTGTATCACC  (R) GGTCAGCATGTTGTACCACAGG |
| ACTB | (F) CAC CAT TGG CAA TGA GCG GTT C  (R) AGG TCT TTG CGG ATG TCC ACG T |
| (F): forward; (R): reverse | |
